# Evolution of ivermectin resistance in the nematode model *Caenorhabditis elegans*: critical influence of population size and unexpected cross-resistance to emodepside

**DOI:** 10.1101/2024.12.03.626540

**Authors:** Jacqueline Hellinga, Barbora Trubenova, Jessica Wagner, Roland R. Regoes, Jürgen Krücken, Hinrich Schulenburg, Georg von Samson-Himmelstjerna

## Abstract

The emergence and spread of anthelmintic resistance represent a major challenge for treating parasitic nematodes, threatening mass-drug control programs in humans and zoonotic species. Currently, experimental evidence to understand the influence of management (e.g., treatment intensity and frequency) and parasite-associated factors (e.g., genetic variation, population size and mutation rates) is lacking. To rectify this knowledge gap, we performed controlled evolution experiments with the model nematode *Caenorhabditis elegans* and further evaluated the evolution dynamics with a computational model. Large population size was critical for rapid ivermectin resistance evolution *in vitro* and *in silico*. Male nematodes were favored during resistance evolution, indicating a selective advantage of sexual recombination under drug pressure *in vitro*. Ivermectin resistance evolution led to the expected emergence of cross-resistance to the structurally related anthelmintic moxidectin but unexpectedly also to the structurally unrelated anthelmintic emodepside that has an entirely different mode of action. In contrast, albendazole, levamisole, and monepantel efficacy were not influenced by the evolution of Ivermectin resistance. We conclude that combining computational modeling with *in vitro* evolution experiments to test specific aspects of evolution directly represents a promising approach to guide the development of novel treatment strategies to anticipate and mitigate resistance evolution in parasitic nematodes.

## Introduction

Ivermectin (IVM) belongs to the anthelmintic class of macrocyclic lactones (MLs) (1). It was first discovered in the late 1970s, and due to its unprecedented spectrum of efficacy and excellent safety profile, it quickly became one of the most often used pharmaceutical compounds in veterinary medicine. Subsequently, IVM was also developed for use in humans, e.g., against various filarial infections, including *Onchocerca volvulus*, the cause of river blindness in Africa (2). This use of IVM eventually led to the award of the Nobel Prize in Physiology or Medicine to its discoverers Satoshi Omura and William C. Campbell in 2015 (3). It is still widely used against several parasitic worms in veterinary and human medicine with several pharmaceutical formulations (4–6).

In nematodes, IVM binds irreversibly to inhibitory glutamate-gated chloride ion (GluCl) channels on membranes of (e.g., pharyngeal) muscles, motor neurons, or the reproductive tract, leading to an influx of chloride ions (5). This causes permanent hyperpolarization of the neuronal membrane and, ultimately, paralysis of the respective muscles (4,5,7–9). The GluCl channel consists of five membrane spanning subunits which are expressed by a species-specific divergent number of genes, for example, in *Caenorhabditis elegans* by six genes – i.e., *avr-14*, *avr-15*, *glc-1*, *glc-2*, *glc-3*, and *glc-4*. The sensitivity of the GluCl channels depends on the respective subunit composition, which has been found to differ between tissues in a given nematode species and between species (5).

Since the introduction of IVM and its widespread use, there have been reports of growing resistance in veterinary and human medicine (10–14). There are multiple pathways thought to cause resistance; however, the exact mechanism of resistance is not yet known, and it may well differ between nematode species and even between different populations of the same species. Dent et al. 2000 reported that mutation at two of the three GluCl genes (*glc-1, avr-14,* and *avr-15*) in *C. elegans* led to resistance (15). Resistance to IVM was reported for *Cooperia oncophora* with a L256F mutation in AVR-14 α-type subunit of a glutamate-gated chloride channel (16). Also, in *C. oncophora,* IVM resistance was seen when AVR-14B was knocked out (17). Morover, P-glycoproteins (Pgps) are likely to contribute to IVM resistance, as their overexpression was observed in IVM resistant *H. contortus* where the sheep were infected from a known IVM-resistant *H. contortus* line (18). Much of the work describing the expression of Pgps in IVM resistance has been done in *C. elegans,* where parasite Pgp genes are transgenically added to *C. elegans* (19,20).

IVM resistance mechanism was further linked to a locus on chromosome 5 of *Haemonchus contortus* (21). This was subsequently confirmed through quantitative trait loci mapping of C. elegans lines, revealing two IVM resistance loci on chromosome 5 (22). The first locus contains one of the previously described genes, *glc-1*, while the second locus has not been described before. Ménez et al. 2019 reported that knockout of the nuclear hormone receptor gene *nhr-8* in *C. elegans* resulted in sensitivity, and strains with deletions in the nhr-8 gene in its DNA-binding domain also showed increased sensitivity to IVM. Also, in this *C. elegans nhr-8* mutation strain, *pgp-6* was overexpressed (23). While many pathways are suspected to be responsible for resistance, the exact mechanism(s) and also their evolutionary emergence remains elusive. Previous studies have concentrated on the genetic and molecular basis of IVM resistance; however, additional research into treatment frequency and intensity, genetic diversity, and reproductive strategies is vital for advancing sustainable parasite control methods.

The free-living non-parasitic nematode *C. elegans* represents a highly informative model for dissecting the molecular targets of many anthelmintics and associated resistance mechanisms. The use of *C. elegans* is appealing as it has a short development cycle of 3.5 days at 20 °C, can easily be maintained on agar, the genome is excellently annotated, and various genetic tools are available. Due to their short development cycle and the ease of manipulating their population genetics, *C. elegans* has been used in evolutionary biology to answer questions about host-pathogen interactions and the impact of outcrossing (24,25). A study by James and Davey (2009) investigated the evolution of IVM resistance by gradually increasing IVM concentrations over multiple generations. They generated IVM-resistant worms over 44 generations (0 to 6.9 nM) and 60 generations (to 11.4 nM). RT-qPCR analysis revealed overexpression of P-glycoproteins (Pgps) and ABC transporters in resistant worms (26). In 2016, Menez et al. used a similar method to generate a moxidectin (MOX)-resistant *C. elegans* strain (5.7 nM tolerance) and confirmed Pgps upregulation (27). Verapamil, a Pgp inhibitor, increased ML susceptibility in these strains. The IVM and MOX-resistant strains from these studies are still used to investigate resistance mechanisms (28,29). Figueiredo et al. (2018) further applied this step-wise resistance evolution approach to develop a *C. elegans* strain resistant to 34.2 nM IVM (30). While trying to uncover the IVM-resistance mechanism, these studies left gaps by not asking specific questions to address how individual factors (i.e., population size, genetic variation, and treatment intensity) contribute to IVM resistance.

Population biological and genetic modeling, while used extensively to understand and predict drug resistance evolution of viruses, bacteria, and eukaryotic microparasites (31), has been less commonly applied to the study of anthelmintic resistance evolution in parasitic metazoans, including nematodes. Most computational models for parasitic worms investigated the role of susceptible strain reservoirs in resistance spread (32–36). These models offer valuable insights into sustainable treatment strategies. Still, the complexities of polygenic resistance and the impact of critical factors like ploidy and reproductive strategies or genetic variation are less frequently addressed in existing models. However, the effect of genetic architecture, dominance, and mating preference on the rate of resistance evolution has already been emphasized (36–38), highlighting the need for new, tailor-made polygenic models of drug resistance evolution.

Our study aims to use *C. elegans* to combine the experimental observations with computational modeling to answer how the intrinsic pathogen factors of population size and genetic variation through sexual recombination affect the rate of ivermectin resistance evolution. The *in vitro* experimental approach presented here relies on previous protocols testing the influence of population size on *C. elegans*-pathogen coevolution (39) and the step-wise introduction of IVM to *C. elegans* (26). The study reported here is the first IVM resistance evolution experiment to include biological replicates and male outcrossing during the evolution process and to report cross-resistance to emodepside. The predictions made by the computational model about the impact of population size and genetic variation with male outcrossing were confirmed by the *in vitro* experiment. Therefore, more computational models based on this work would allow for the simulation of complex scenarios and factors that might be difficult or costly to manipulate directly in a laboratory setting and will enable us to simulate the experiment with double duration, providing additional insight into the long-term consequences of the investigated factors. Bringing in the ability to answer the perplexing field related questions that have remained unanswered by the leading anthelmintic resistant research groups.

## Materials and methods

### Worm strain maintenance and Escherichia coli preparation

*Caenorhabditis elegans* strain WBM1133 (wbmIs63 [myo-3p::3XFLAG::wrmScarlet::unc-54 3’UTR *wbmIs61]) (40) was obtained from the Caenorhabditis Genetics Center (CGC). The strain expresses a red fluorescent protein in its body wall muscles and was described to have no obvious phenotype (40). Worms were maintained on nematode growth medium (NGM) agar supplemented with *E. coli* OP50 as a food source at 16 °C using standard manipulation methods (41). Males were introduced into the population by seeding an NGM plate with 10 fourth stage larvae (L4) that were heat shocked at 35 °C for 2 h. The male population was maintained by picking at a propagation ratio of 3:1 (male: hermaphrodite) at 20 °C. *Escherichia coli* OP50 was prepared as described in Hellinga et. al., (2024) (42).

### Ethyl methanesulfonate (EMS) treatment

To increase genetic variation in our *C. elegans* strain to allow for differences in how resistance is acquired, treatment by ethyl methanesulfonate (EMS) was done, as previously stated in Brenner 1974 (43). The WBM1133 worms containing males were synchronized using bleach, grown to an L4 stage, and resuspended in M9 buffer. Worms were incubated in 50 mM of EMS dissolved in M9 under constant shaking at 20 °C for 4 h. Worms were collected by centrifugation and washed five times with 15 ml of M9. An F2 screen was set up to assess the reproductive health of the EMS-treated worms. Ten gravid worms from EMS-treated and non-treated populations were placed on NGM agar with one worm per plate. The gravid worms were removed after 24 h, and the resulting offspring were counted (Fig. S1).

### Pre-adaptation to Viscous medium

The experimental evolution was performed in a Viscous medium (V medium) (39,42). Therefore, pre-adaptation of the *C. elegans* to V medium was done to ensure the worm populations were adapted to the main conditions of the experimental approach, and the V medium did not act as an additional selective factor during the main experiment (39). In the pre-adaptation process 3.2 ml of Viscous medium (S basal (10 mM potassium phosphate buffer pH 6.0, 100 mM NaCl), 0.8% hydroxypropyl methylcellulose, 10 mM potassium citrate pH 6.0, 3 mM Mg_2_SO_4_, 3 mM CaCl_2_, 5 µg/ml cholesterol) was put into 6-well culture plate (Sarstedt #83.3920) supplemented with *E. coli* OP50 (final optical density, OD_600_ = 2.0). Worms were incubated at 20 °C with constant shaking at 150 rpm for 3-4 days. The worms were pooled at each transfer and washed with 200× volume of S basal. The size of the worm population was counted and set to 3000 worms per well. In total, 15 transfers were done; at transfers 1, 5, 7, 8, 9, 10, and 12, a male-rich immigrant population was added as 10% of the total population to maintain the percentage of males during the adaptation at 30%. To add further genetic variation, at transfer 2, male induced worms from H. Schulenburg laboratory were added to the population corresponding to 16% of the total worm population. For each transfer, 15 drops (10 µl each) were mixed with 1 µl 5% Lugol’s iodine to count the number of males in the population. Only L4 and older worms were counted, and the worm was determined to be a male if it had either the characteristic male tail or lacked 2 gonad arms or eggs. After the 15^th^ transfer, worms from 24 6-well plates were pooled and divided into 8 batches containing 22% males. These populations were synchronized by bleaching, and the corresponding L1 worms were pooled and used in the evolutionary experiment.

### In vitro evolution experiment

Synchronized L1 worms from the above were divided into 72 biological replicates for the evolutionary experiment. Half of the populations (n = 36) were used for selection with IVM, and the other half was used as a control for genetic drift in the absence of IVM. For each of these two lines, 12 replicates were used for each population size. The population sizes were set to 200, 1000, and 2000 worms. The control populations were exposed to 0.1% DMSO for the evolution experiment. The IVM-treated populations were exposed to IVM in a step-wise manner. Ivermectin concentrations started with 0.1 nM and were doubled until 1.0 nM; all following concentration steps were 25 % - 33 % higher than the last. All populations were grown in 12-well culture plates (Sarstedt #83.3920); each well contained 1.6 ml Viscous medium supplemented with *E. coli* OP50 (final optical density OD_600_ 1.3 for 2000 worms, 1.0 for 1000 worms, and 0.5 for 200 worms). Worms were incubated at 20 °C with constant shaking at 150 rpm for four days. On the fourth day, the worm lines were washed with S basal, and 10 drops (5 µl each) were counted. All lines were reset to the original population size and again incubated at 20 °C. The percentage of males was determined for each population replicate as described above. To further protect the evolution experiment, two groups of IVM-treated worms were grown in parallel, with one group of the IVM-treated worms exposed to the introduced concentration of IVM (working group) and the other exposed to a greater concentration of IVM (challenge group) (Fig S2). Once all challenge groups acquired resistance to a specific concentration of IVM, they were kept at that concentration for one more generation so each biological replicate could be expanded to freeze multiple vials for future analysis. The challenge group then became the working group, and a new challenge group was established (Fig. S2). If resistance was not acquired for the challenge group, then the challenge group was discarded, and propagation continued from the working group. A population was only stated to have acquired resistance to the respective IVM concentration once all 12 biological replicates produced a population size 10 times higher than the starting population, and the worms were seen to move freely (Fig. S3). Finally, the 1000 worm populations were grown for 38 generations, and the 2000 and 200 worm populations were grown for 39 generations.

### In silico evolution experiment

The computational model used to simulate the evolution experiment was based on the population genetic, compartmental model proposed in Trubenová et al., (2024) (44) with a few modifications, as explained in the Supplementary material. The model considered IVM resistance a polygenic trait, with 6 independent diploid loci (genes) of various effects on the level of resistance. This generated 729 possible genotypes, providing substantial complexity to reflect the diverse resistant strains observed in experimental settings while remaining computationally tractable (Fig. S4). Resistance mutations were recessive, meaning both copies of a gene had to be mutated for resistance to increase. In addition, a trade-off was incorporated: mutations increased IVM resistance but assigned a lower fitness to the worms in a drug-free environment. We used empirically-supported pharmacodynamic relationships between the worm fitness (measured as egg count) and the ivermectin concentration to determine the fitness of various genotypes under increasing ivermectin concentrations (see Supplementary material, Fig S4, Table S1).

The simulations mirrored the steps of the *in vitro* experiment (explained above), starting with susceptible *C. elegans* populations of different sizes (200, 1000, 2000) containing 20% males. The simulations of the evolutionary process were then run for a set number of cycles, corresponding to generations. In each generation, hermaphrodites self-fertilized or mated, generating new genotypic combinations assuming no linkage among the loci. Mutations occurred at a small rate (0.0001 mutations per genome) (Table S2), generating new variants.

If the population grew above a given threshold (30-fold increase), it was considered adapted, and the IVM concentration was increased (Table S3), mimicking the experimental protocol. At the end of each cycle, populations were diluted to the original size. Simulations were stochastic and repeated 100 times for each population size over 40 generations following the process outlined earlier. In addition, the same evolutionary scenario was modeled for a longer time of 80 generations, with stochastic simulations repeated 50 times for each population size. See Supplementary material for a detailed description of the simulation process and simulated scenarios.

### Larval development assays

Frozen ancestor and endpoint (generations 38/39) evolved worms were thawed and regenerated for two generations without drug pressure in V medium. Adults from the F2 generation were synchronized by bleaching. The L1 worms were counted, and approximately 100 worms were added in 24 µl to every well of a 48-well microtiter plate already containing 200 µl V medium, 25 µl *E. coli* OP50 (OD_600_ =2), and 1 µl anthelmintic. Albendazole (ABZ), emodepside (EMO), IVM, monepantel (MON), and moxidectin (MOX) were dissolved in 100% DMSO, while levamisole (LEV) was dissolved in S basal. Ivermectin was added in a serial dilution to the final concentration range of 0-8 nM with a 1.4-fold dilution for the control and ancestor populations and 0-110 nM with a 1.4-fold dilution for the resistant populations. For MOX, a 1.5-fold dilution was done with the final concentration range of 0-6 nM for the control and ancestor populations and 0-75 nM for the resistant populations. Emodepside had a 1.4-fold dilution with a final concentration range of 0-100 nM for the control and ancestor populations and 0-600 nM for the resistant populations. Albendazole had a final concentration range of 0-100 µM with a 1.5-fold dilution, MON had a final range of 0-100 nM with a 1.4-fold dilution, and LEV had a final range of 0-250 µM with a 1.6-fold dilution. Worms were incubated at 20 °C with constant shaking at 150 rpm. After a 48-52 h incubation, the development was stopped by adding Lugol’s iodine. Developed worms were either L4 or adult stage (males and hermaphrodites), while non-developed worms were L1, L2, and L3. For each resistant/control population, a biological replicate of the assay was performed once with anthelmintic drug concentrations in triplicate. The ancestor population was used as a control in all experiments. The mean EC_50_ values were determined by fitting a variable slope log ([IVM]) vs. response (percentage developed worms) four parameters logit curve using GraphPad Prism 5.03.

### Statistical analysis of the larval development assay and the in vitro experiment

Standard deviations and significant differences between the mean EC_50_ of the larval development assay were determined by the extra sum of squares F-test within GraphPad Prism. P values < 0.05 were considered to be statistically significant. For development analysis, significant differences in development between the ancestor and the endpoint evolved lines were calculated using a non-parametric Kruskal-Wallis test followed by a Dunn’s post-hoc test comparing the evolved lines to the ancestor. ANOVA was used to determine significant differences in the percentage of males during the *in vitro* evolution experiment, followed by a Tukey’s Multiple Comparison posthoc test in GraphPad Prism.

## Results

### Preparation of worms for in vitro evolution experiment

The study investigated how population size impacts the rate at which *C. elegans* acquire IVM resistance (Fig. 1A). Consistent with previous experiments on IVM resistance in *C. elegans,* a hermaphrodite N2 strain was sought after (26,27). The WBM1133 strain was selected for its characteristic *mScarlet* gene under a *myo-3* promoter on chromosome 1 (40) for future work where tracking the resistant population by fluorescence would be necessary. Additionally, males were incorporated into the *C. elegans* strain to minimize differences in the number of deleterious mutations and increase genetic variation across biological replicates and population sizes. This inclusion of males facilitated outcrossing events, leading to a single lineage with adaptive mutations instead of multiple selfing lineages with diverse adaptive mutations. The WBM1133 strain maintained males over several generations, unlike the WT N2 strain. Once the male frequency in the WBM1133 population stabilized, EMS treatment was performed as described by Brenner et al. (1974) (43). After treatment, the F1 generation’s progeny count was similar to that of non-EMS-treated worms (Fig. S1). Visual monitoring showed no physical defects in body width, brood size, or rolling.

**Fig. 1.**
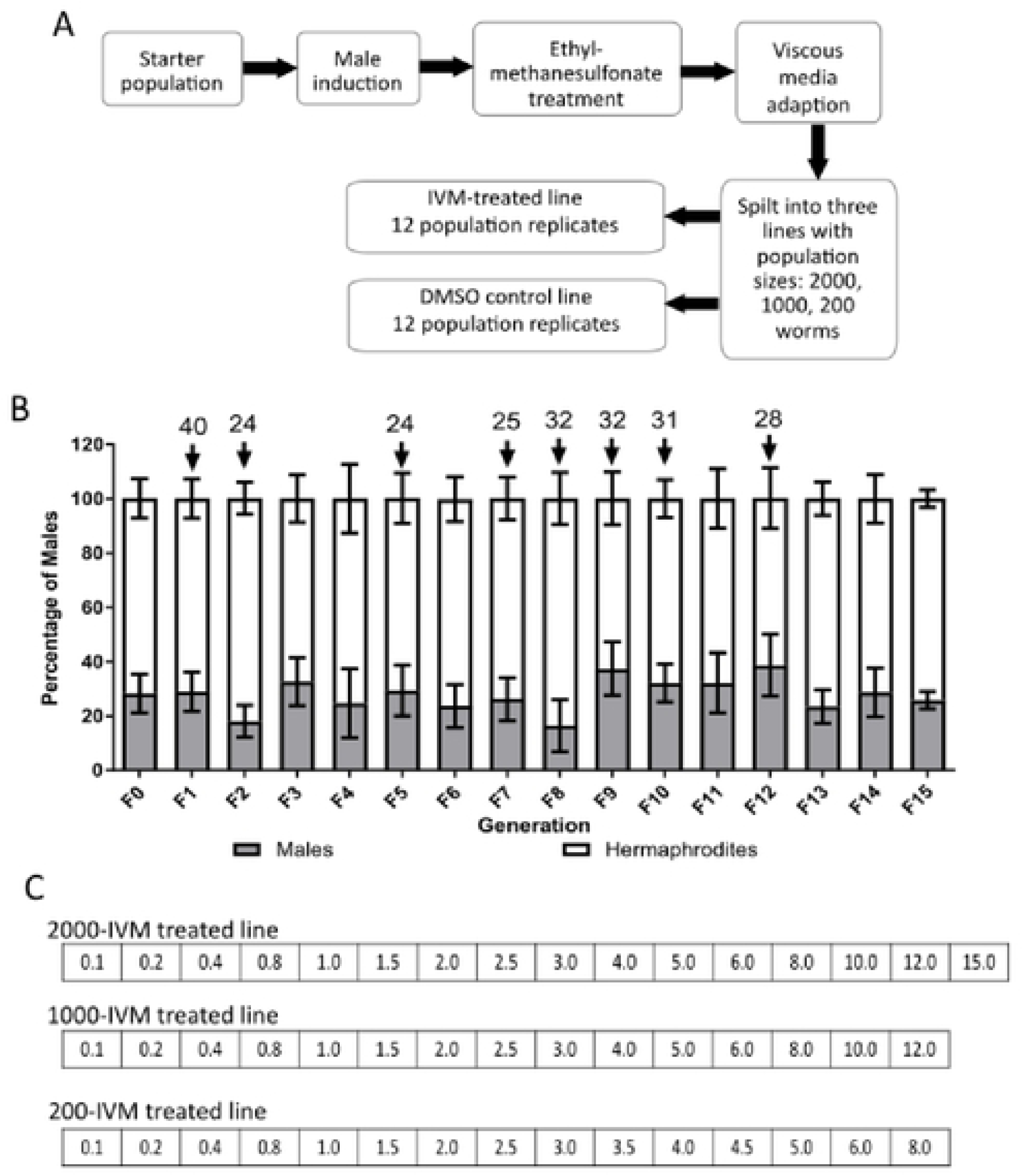
Setup of the *in vitro* evolution experiment with *C.elegans.* (A) Workflow of in vitro evolution experiment including V medium pre-adaptation and the start of the evolution experiment. (B) Percentage of males in each generation during the V medium pre-adaptation. The arrows indicate when immigrant males from NGM agar plates were added. The numbers above the arrows indicate the percentage of males in the immigrant population. Data show means± standard deviation for hermaphrodite and male percentages. (C) Concentrations ofIVM that were used for the in vitro experiment for the three population lines.

The EMS-treated population was placed in V medium for 15 generations to adapt before IVM selection began. The adaptation to the V medium was performed to ensure the worms were acclimated to the primary conditions of the experimental approach and to prevent the V medium from acting as an additional selective factor during the main experiment. Immigrant males were added during this period to maintain male frequency (Fig 1B). The male reduction likely resulted from the absence of a stressor requiring males to adapt or from the high progeny produced by selfing. When male percentages in the V medium fell below 25%, 10% of the population was supplemented with males from NGM agar plates, where male percentages were above 30%. No immigrant males were added during the last three generations to avoid influencing the adaptation.

### Population size increases the rate of ivermectin resistance evolution during the in vitro evolution experiment

For the *in vitro* evolution experiment, worms adapted to V medium were synchronized using the standard bleaching method and divided into three population sizes: 200, 1000, and 2000 worms, with 12 biological replicates for both control and IVM-treated lines. Worms were exposed to increasing IVM concentrations: 0.1 nM for one generation, 0.2 nM for one generation, and 0.4 nM for two generations. When introduced to 0.8 nM IVM, the 200-worm populations took three generations to evolve resistance, while the 1000 and 2000 populations took only two generations. Throughout the experiment, the 200 IVM-treated line took 3 generations to adapt to a given IVM concentration. The 1000 IVM-treated line also took 3 generations but faced a larger IVM concentration step. The 2000 IVM-treated line generally adapted within 2 generations. However, at one point during the evolution experiment, each IVM-treated line required more than four generations to adapt to the next high IVM concentration. For example, this was observed for the 2000 IVM-treated line when it was introduced to 6.0 nM IVM. Instead of taking two generations to adapt, it took 6 generations. The 1000 IVM-treated line required 5 generations at 5.0 nM IVM, and the 200 IVM-treated line required 4 generations at a concentration of 2.0 nM. Lastly, smaller IVM increments were used for the 200 IVM-treated line to prevent extinction. The experiment was initially planned for 40 generations but was concluded early when the 1000 IVM-treated line adapted to 12 nM IVM by generation 38, and the 2000 and 200 IVM-treated lines adapted to 15 nM and 8 nM IVM, respectively, by generation 39 (Fig. 2A). The results demonstrated that population size influenced the rate of IVM resistance evolution.

**Fig. 2.**
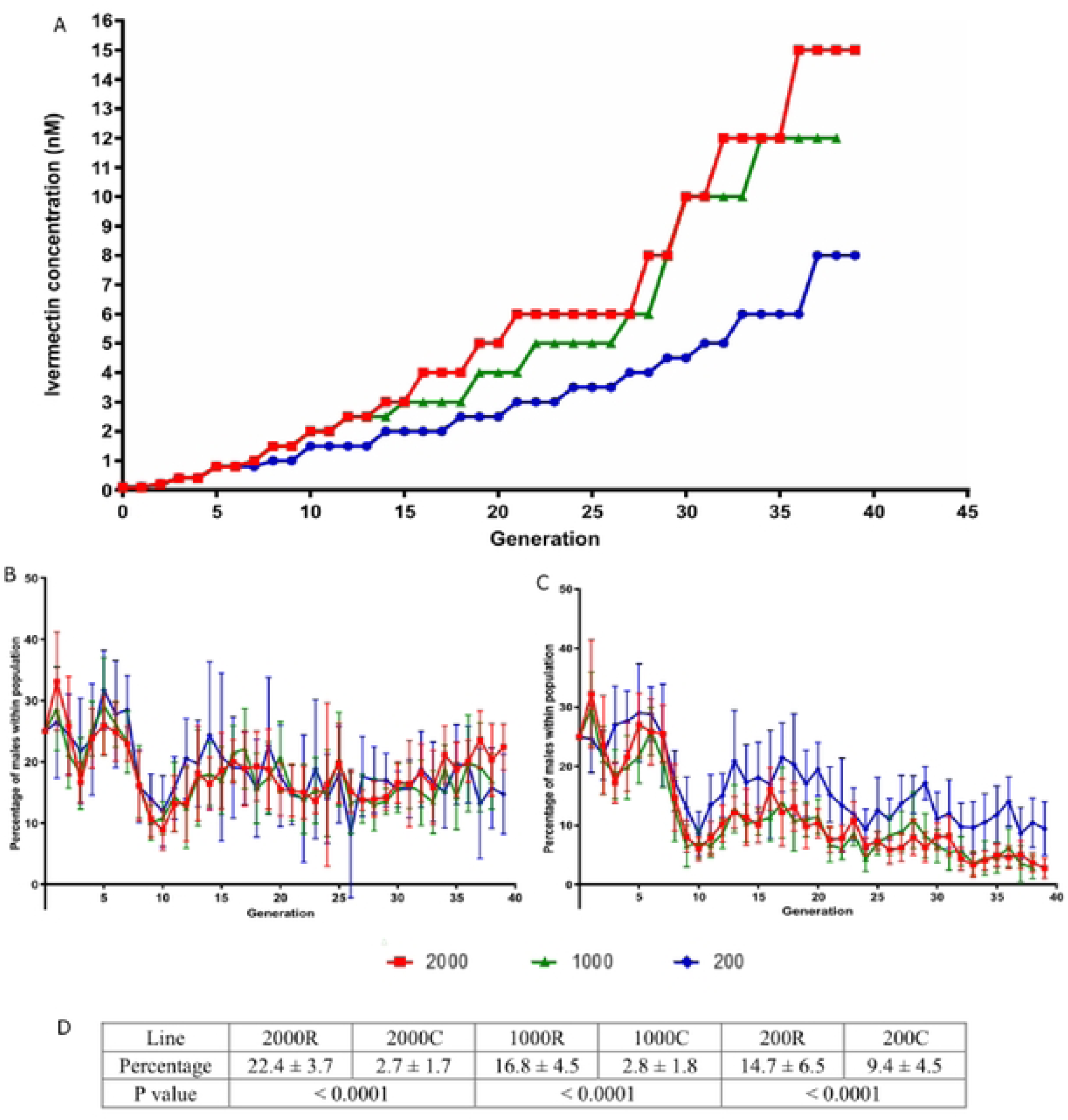
Outcome of the *in vitro* evolution experiment. Population sizes are indicated by red squares (2000), green triangles (1000) and blue circles (200). (A) Concentration ofIVM tolerated by *C. elegans* populations during the *in vitro* evolution experiment. (B, C) Percentage of males found in IVM-treated lines (B) and control lines (C). Graphs show the mean and the standard deviation of the 12 biological replicates,. Adult stagc/L4 stage worms were counted in 10 10 µl drops. Worms were paralyzed ,vith Lugol’s reagent so that anatomical structures could be observed. (D) Percentage of males observed in the endpoint generation of the *in vitro* evolution experiment. P values were determined by one-way ANOVA and Tukey’s multiple comparison post-test comparing the percentage of males between the resistant and control populations.

### Ivermectin selection led to a significantly higher frequency of males in all population sizes

Initially, males were introduced to the *C. elegans* population to ensure a possibility for outcrossing as a basis for maintaining genetic variation and to minimize the number of deleterious mutations (45) within the biological replicates for each population size. During the adaptation process to V medium, the number of males gradually decreased over time and had to be supplemented with immigrant males. However, in the IVM-treated lines, males were easily maintained throughout the evolution experiment. In the first 10 generations, the percentage of males was similar in both control and IVM-treated lines (Fig. 2B, C). Between generations 5 and 10, there was an overall decrease in males across all populations. At generation 11, the percentage of males in the IVM-treated lines rebounded to approximately 20% and remained stable for the rest of the experiment, while the percentage in the control lines slightly rebounded but overall decreased over time (Fig. 2B, C).

Interestingly, the percentage of males in the final generation of the IVM-treated lines decreased with smaller population sizes: the 2000 IVM-treated line had the highest percentage of males at 22.4%, the 1000 IVM-treated line had 16.8%, and the 200 IVM-treated line had the lowest at 14.7% (Fig. 2D).

The differences in the percentage of males between IVM-treated and control populations were statistically significant (P value < 0.001, One way ANOVA, Tukey’s Multiple Comparison Test) (Fig. 2D). The higher percentage of males in the 200 control line compared to the 1000 and 2000 control lines could be an artifact due to fewer worms in the L4/adult stage being counted. Overall, the observed percentage of males in this evolution experiment aligns with previous findings where environmental stressors, such as different pathogenic bacteria, increased the number of outcrossing events (46) or ensured the maintenance of males (47).

### Computational modeling shows the impact of population size on both evolutionary rate and trajectory

We developed a computational model to investigate if the experimental observations are consistent qualitatively and quantitatively with the population biology and genetic characteristics of the system, such as population sizes, the mutation rate of *C. elegans*, and its pharmacodynamics. In a nutshell, the model describes the reproduction and evolution of the worms in the experiment, assuming six diploid resistance loci (see Materials & Methods). The computational model in this experiment was previously used to understand the role of different reproduction methods in resistance evolution to anthelmintics (44).

As in the *in vitro* evolution experiment, our model shows a clear advantage for larger populations in the rate of resistance evolution. This reflects the experimental finding that the 2000 population size outpaced the 1000 and 200 population sizes in adapting to increasing IVM concentrations. In addition, the simulations revealed several important insights into the evolution of IVM resistance: Firstly, we observed that all populations adapted to low IVM concentrations (up to 2.0 nM) within the first nine generations. The EC_50_ of the computational model for the initial population was set to 1 nM to reflect literature values for wild-type *C. elegans*. Therefore, even the wild-type worms can reproduce sufficiently at these concentrations, increasing the average egg count above 30 (Fig. 3A). However, individual simulations showed varying outcomes at higher concentrations. While some populations failed to adapt beyond 2.0 nM, others, even those with small population sizes, adapted to 15 nM within 40 generations. This highlights the impact of random mutations, which become especially significant in smaller populations. Larger populations showed more consistent adaptation trajectories, as they had a higher supply of beneficial mutations (Fig. 3B, C). In agreement with experimental observations, the mean (Fig. 4A) and distribution (Fig. 3D, E) of the final IVM concentrations reached different levels between population sizes.

**Fig. 3.**
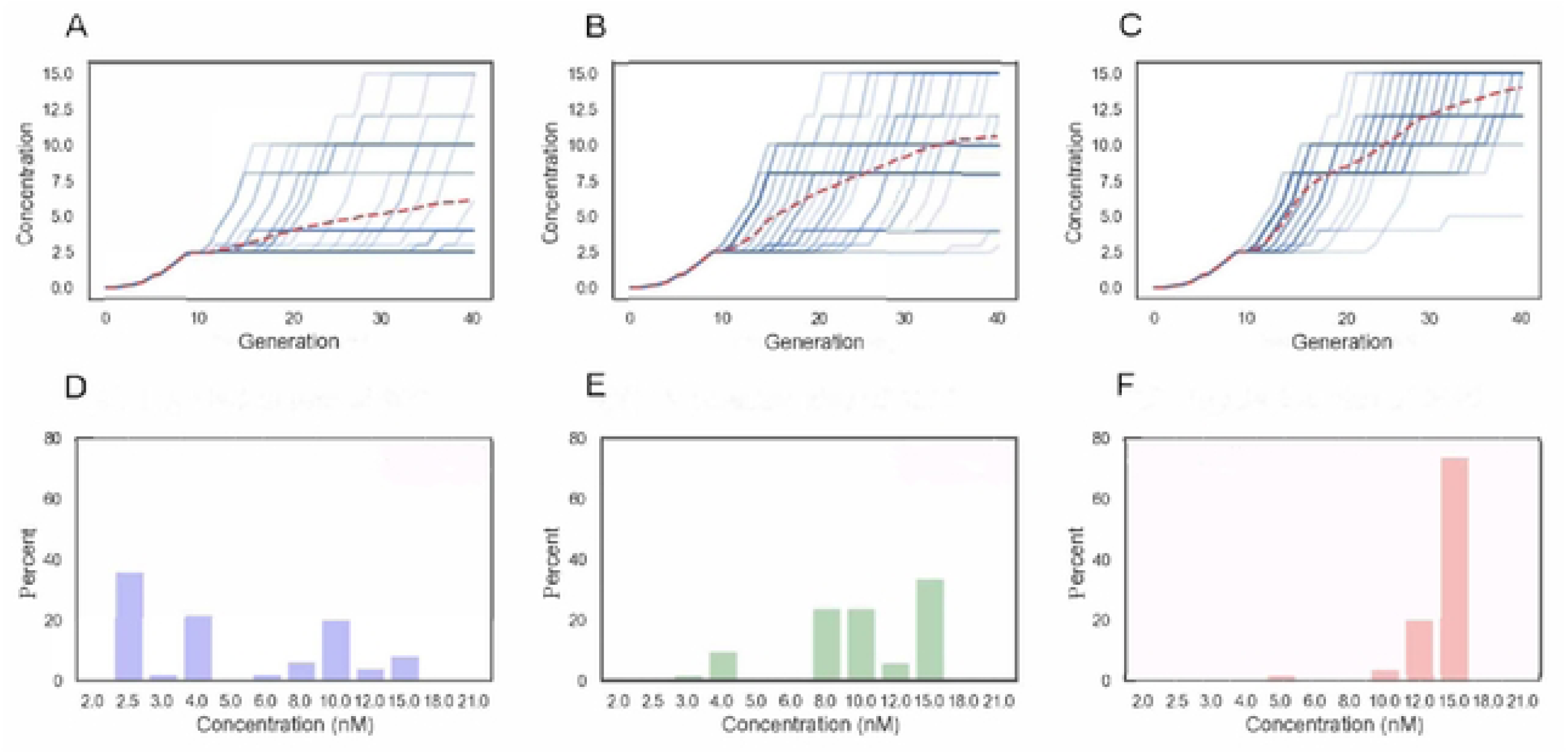
Computational model of three *Caenorhabdilis elegans* population sizes evolving to ivennectin over 40 generations. The concentration at which the worm population grew as a function of time in the simulated evolution experiments (A-C). Blue solid lines correspond to individual simulations (n = 100); the red dashed lines represent the means. Distribution of the final ivennectin concentrations of the *C. elegans* populations adapted after 40 generations (0-F). A, D: population size 200. B, E) population size 1000, C, F) population size 2000.

**Fig. 4.**
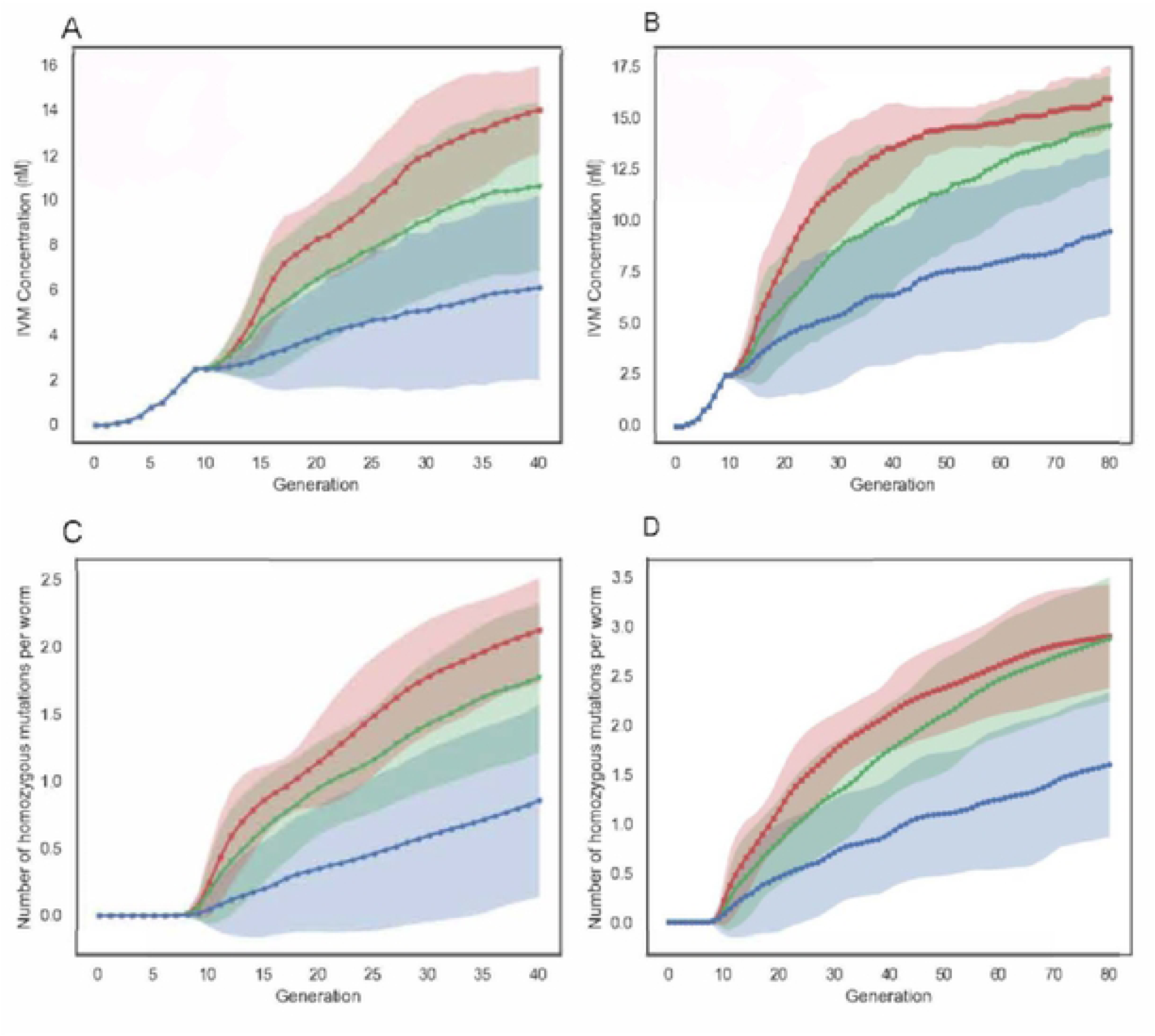
lvenncctin concentration range the *Caenorhabditis elegans* populations adapted to over time and the number of homozygous mutations predicted to occur while adapting. The 200 population are shown in blue, 1000 in green, and 2000 in red. (A, B) Modeling results for concentrations at wh.ich the wonn populations arc cultured as a function of time. (A) 40 generations and (B) 80 generations. The line represents the mean, and the standard deviations are the shaded areas. (C, **0)** The average number of homozygous mutations per wonn present within the population at a certain time point: (C) 40 generations and (0) 80 generations, mean (lines) and standard deviation (shaded areas).

In the extended simulations of 80 generations, smaller populations (200 and 1000) adapted to higher IVM concentrations (Fig. 4B). Interestingly, while the largest population (2000) also reached, on average, resistance to a higher concentration, its median adaptation (15 nM) remained similar to the smaller populations (15 nM for population of 1000 worms, 10 nM for populations of 200 worms, Fig. S7). This suggests that while initial adaptation is more likely in larger populations due to increased mutation supply, the difference between 1000 and 2000 individuals may not be large enough to impact long-term adaptation rates significantly. While the average number of homozygous mutations increased over time in all population sizes (Fig. 4C, D), as stated above, none of the populations reached a state in which all loci would be mutated and homozygous, indicating the potential for further evolution, even after 80 generations.

To deepen our insight into the outcome of the evolutionary process, we investigated the genetic composition of the final populations after 40 generations of evolution. Our simulations revealed substantial variation in the genetic makeup of resistant populations. Population size had an impact on which resistance loci were fixed in the population (Fig. 5). In small populations (200 individuals), any resistance mutation had a similar chance of becoming fixed, likely due to the limited mutation supply (Fig. 5A). However, in larger populations (1000 and 2000), mutations with larger effects were more likely to dominate (Fig. 5B, C). This suggests that clonal interference (competition between strains with different beneficial mutations), rather than mutational supply, plays a role in these larger populations.

**Fig. 5.**
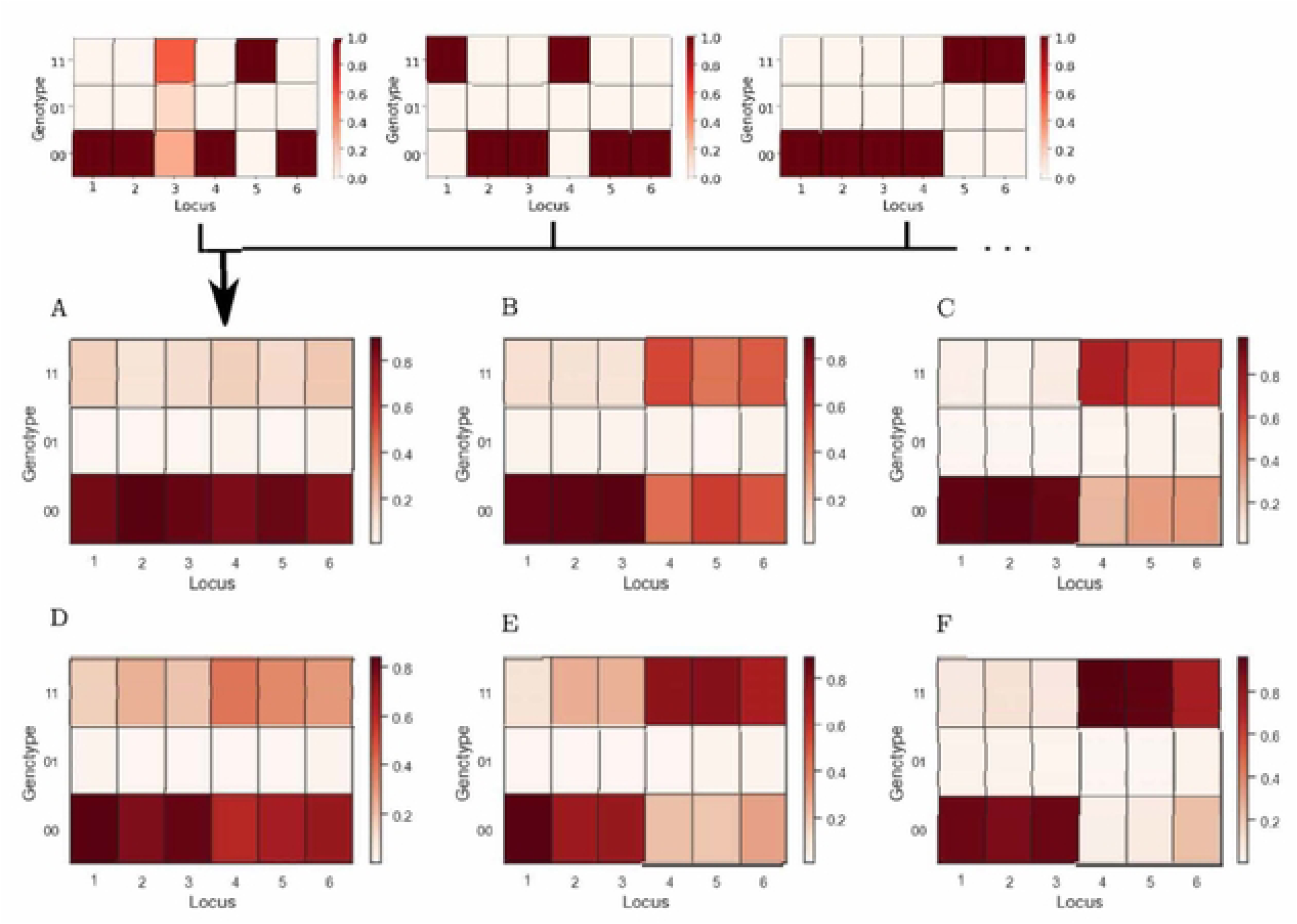
Genetic composition after 40 generations of evolution modeling using 100 individual simulations. Different loci are portrayed on the **X** axis, and on the **Y** axis the possible genotypes for each locus. The rectangle’s color corresponds to the fraction of individuals having a particular genotype on that particular locus. Values in each column add to 1. **(A-C)** Genetic composition of the final populations reached after 40 generations of evolution. (D-F) Genetic composition of the final populations reached after 80 generations of evolution. **(A, D)** 200 populations, (B, E) 1000 populations, (C, F) 2000 populations.

In populations that evolved over 80 generations, a clear preference for mutations with larger effects emerged over time, even in the smallest populations (Fig. 5 D-F). Additionally, in the largest population, simulations revealed selection against mutations with the highest cost (locus 6), favoring those with equal benefits but lower costs (loci 4 and 5) (Fig. 5F). Combining this insight with the previous observations, we hypothesize that while the mutations of three large-effect loci were sufficient to allow populations to adapt to 15 nM, the beneficial effects of the remaining three loci were too weak to allow further adaptation. In addition, fitness costs accumulated, preventing further adaptation.

### Emergence of cross-resistance of the in vitro ivermectin-adapted populations to the macrocyclic lactones ivermectin and moxidectin

The final generation of each biological replicate from all IVM-treated and control lines was characterized phenotypically with the help of larval development assays to assess the exact level of evolved IVM resistance across the evolution experiment. Ancestor populations were also tested, i.e., the 15^th^ generation of the pre-adaptation process to the V medium. All twelve biological replicates from IVM-treated and control lines were tested for each population size.

The 2000 IVM-treated line had a mean EC_50_ value of 3.48 nM, which is 9.40 times higher than the ancestor’s EC_50_ value of 0.37 nM (Table 1; Fig. 6A). The 1000 IVM-treated line had a mean EC_50_ value of 3.21 nM, 8.67 times higher than the ancestor (Table 1). The 200 IVM-treated line had a mean EC_50_ value of 2.88 nM, 7.78 times higher than the ancestor (Table 1). Thus, all IVM-treated lines were significantly more resistant to IVM than the ancestor (P value < 0.0001). The fold change of resistance compared to the ancestor decreased as population size decreased. The control lines of the evolution experiment were also assessed for their resistance level to IVM. The 2000, 1000, and 200 control lines had mean EC_50_ values of 0.36 nM, 0.39 nM, and 0.37 nM, respectively (Table 1). These mean EC_50_ values were not statistically different from the ancestor’s EC_50_ values (P values: 0.3337 for 2000, 0.336 for 1000, and 0.860 for 200). This suggests that the conditions of the evolution experiment did not favor resistance evolution and that resistance was derived from the step-wise introduction of IVM (Fig. 6A). All population sizes for each IVM-treated and control lines had similar EC_50_ values, indicating no outlier biological replicate.

**Fig. 6.**
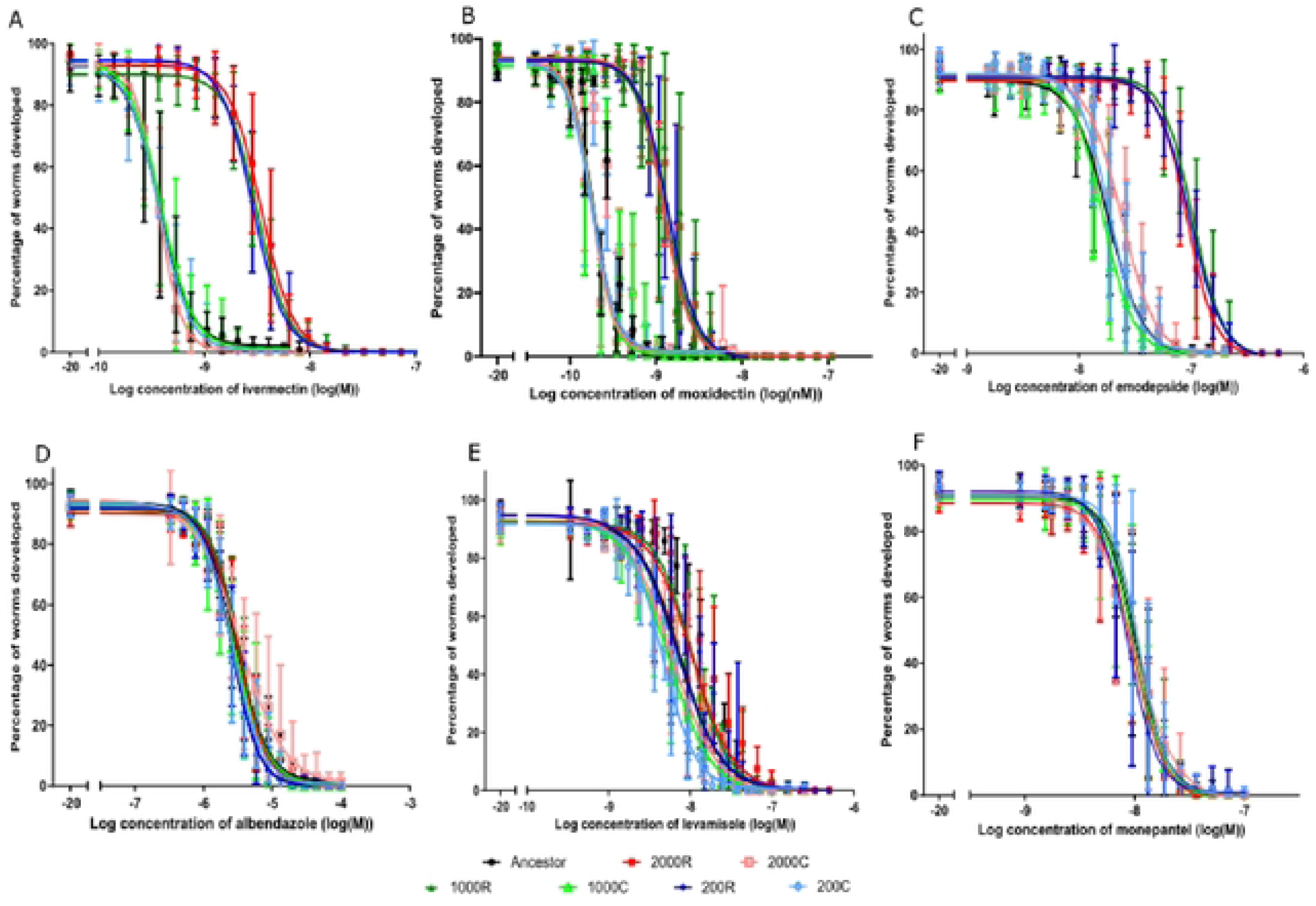
Concentration response curves profiling the responses of the ancestor and the endpoint of the IVM-treated and control lines in an *in vitro* larval development assay. Ancestor is black square, the 2000 IVM-treated line (2000R) are shown as red squares, the 2000 control line (2000C) as salmon squares, the 1000 IVM-trcated line (IOOOR) as dark green triangles, the 1000 control line (IOOOC) as light green triangles, the 200 IVM-treated line as dark blue diamonds, and the 200 control line as light blue diamonds. (A) Ivermectin, (B) mox.idectin, (C) emodepside, (D) albendazole, (E) levamisole, (F) monepantel. Plotted are the mean and standard deviation of the 12 population replicates for each worm line. One biological and three technical replicates were done for each population replicate.

**Table 1.** Mean EC_50_ values of the ivermectin resistant lines to macrocyclic lactones, ivermectin, and moxidectin, as determined by larval development assays.

| Anthelmintic | Population | EC<sub>50</sub> [nM] | 95% CI | R<sup>2</sup> | Fold change<sup>a</sup> | P value<sup>b</sup> |
| --- | --- | --- | --- | --- | --- | --- |
| **Ivermectin** | Ancestor | 0.37 | 0.35 – 0.39 | 0.910 | 1.00 |  |
|  | 2000R | 3.48 | 3.34 – 3.63 | 0.938 | 9.40 | < 0.0001 |
|  | 2000C | 0.36 | 0.35 – 0.37 | 0.943 | 0.97 | 0.337 |
|  | 1000R | 3.21 | 3.10 – 3.32 | 0.953 | 8.67 | < 0.0001 |
|  | 1000C | 0.39 | 0.37 – 0.41 | 0.893 | 1.05 | 0.336 |
|  | 200R | 2.88 | 2.78 – 2.99 | 0.953 | 7.78 | < 0.0001 |
|  | 200C | 0.37 | 0.35 – 0.39 | 0.917 | 1.00 | 0.860 |
| **Moxidectin** | Ancestor | 0.19 | 0.18 – 0.19 | 0.963 | 1.00 |  |
|  | 2000R | 1.23 | 1.17 – 1.28 | 0.930 | 6.47 | < 0.0001 |
|  | 2000C | 0.17 | 0.17 – 0.18 | 0.928 | 1.00 | 0.010 |
|  | 1000R | 1.25 | 1.18 – 1.32 | 0.905 | 6.57 | < 0.0001 |
|  | 1000C | 0.18 | 0.17 – 0.19 | 0.910 | 0.94 | 0.127 |
|  | 200R | 1.37 | 1.30 – 1.45 | 0.923 | 7.21 | < 0.0001 |
|  | 200C | 0.18 | 0.17 – 0.19 | 0.901 | 0.94 | 0.060 |

<sup>a</sup>Calculated as EC<sub>50</sub>resistant or control populations/ EC<sub>50</sub> ancestor
<sup>b</sup>Sum of squares F test comparing resistant or sensitive populations to ancestor

The ancestor and the endpoint of the IVM-treated and control lines were also tested against moxidectin (MOX), another macrocyclic lactone, to assess side resistance. The ancestor population had an EC_50_ value of 0.19 nM for moxidectin (Table 1). The 2000 IVM-treated line had a mean EC_50_ value of 1.23 nM, 6.47 times higher than the ancestor (Table 1, Fig. 6B). The 1000 IVM-treated line had a mean EC_50_ value of 1.25 nM, which was 6.57 times higher than the ancestor. Further, the 200 IVM-treated line was the most resistant IVM-treated line to MOX with a mean EC_50_ value of 1.37 nM, which was 7.21 times higher than that of the ancestor. All IVM-treated lines were significantly more resistant to moxidectin than the ancestor (P value < 0.0001 for all population sizes). Remarkably, the fold change in resistance for MOX compared to the ancestor showed an inverse trend to the IVM assays: the 200 IVM-treated line had the highest fold change, and the 2000 IVM-treated line had the lowest. The control lines had mean EC_50_ values of 0.18 nM for the 1000 and 200 lines (Table 1). The 2000 control line (EC_50_ of 0.17 nM) was slightly more sensitive to moxidectin than the ancestor (P value of 0.010), while the other control lines were not significantly different. Finally, all population replicates for the control and IVM-treated lines had similar EC_50_ values (Table 1), confirming no outlier biological replicate.

### Emergence of cross-resistance of the in vitro IVM-adapted populations to emodepside and sensitivity to other anthelmintic classes

The evolved populations were further tested against emodepside, a cyclooctadepsipeptide. The ancestor had an EC_50_ value of 18.27 nM (Table 2). The 2000 IVM-treated line had a mean EC_50_ value of 91.91 nM, 5.03 times higher than the ancestor (Table 2). The 1000 IVM-treated line had a mean EC_50_ value of 103.4 nM, 5.65 times higher than the ancestor (Table 2). Lastly, the 200 IVM-treated line had a mean EC_50_ value of 97.49 nM, 5.33 times higher than the ancestor (Table 2). Therefore, all the IVM-treated lines, which also evolved resistance to moxidectin, showed comparable resistance to emodepside (Fig. 6C). This cross-resistance to emodepside was statistically significant (P value < 0.0001, sum of squares F test). Unlike the trend observed for IVM, resistance to emodepside did not show decreasing EC_50_ values with decreasing population sizes. For the 2000 and 1000 control lines, slightly but significantly higher EC_50_ values (2000: <0.0001, 1000: 0.0006) were observed compared to the ancestor, while no significant difference was detected for the 200 control line (Table 2). No outlier replicate was observed as all population replicates for each IVM-treated and control lines had similar EC_50_ values (Table 2). Therefore, all the IVM-treated lines also evolved resistance to moxidectin and the unrelated anthelmintic emodepside.

**Table 2.**
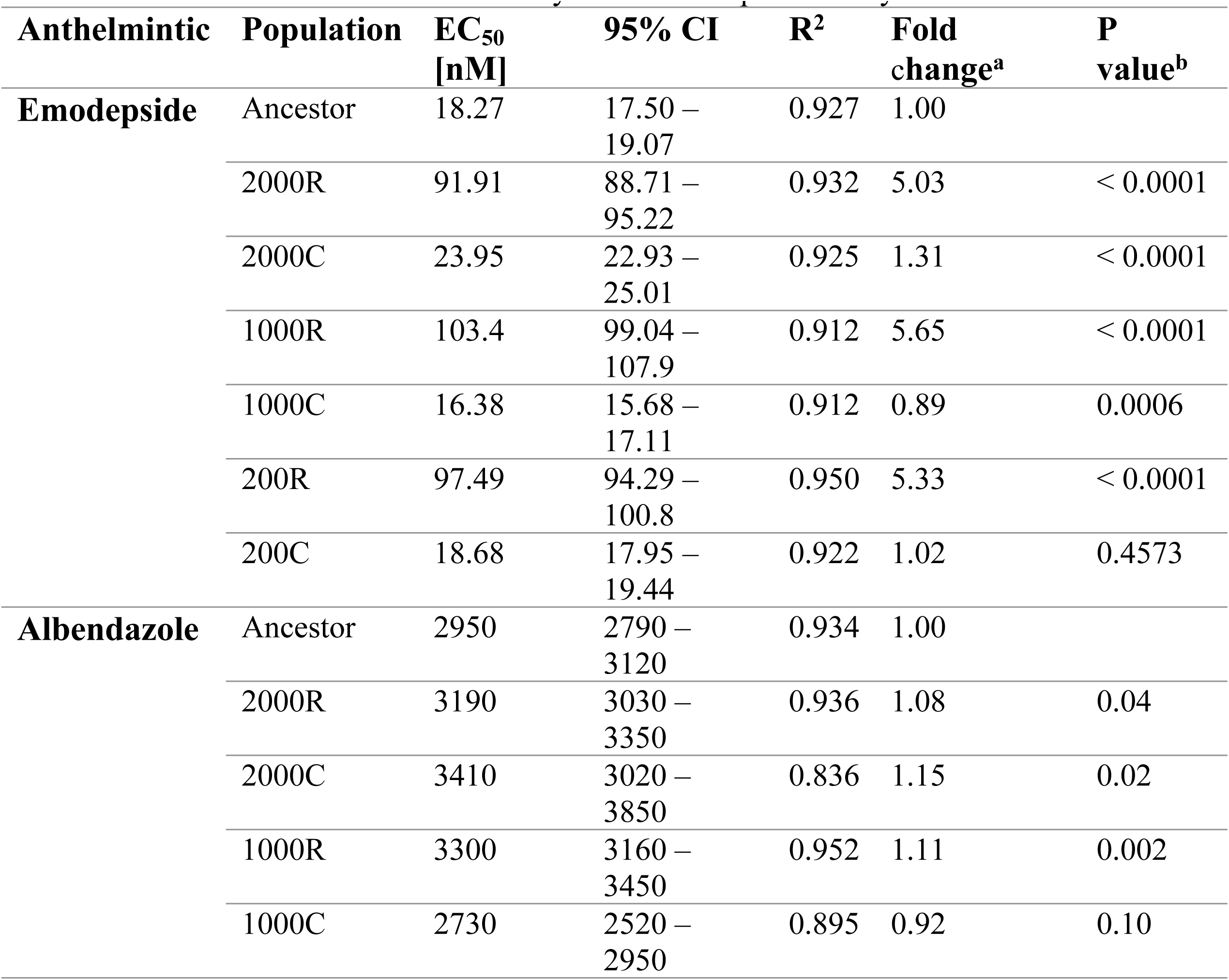
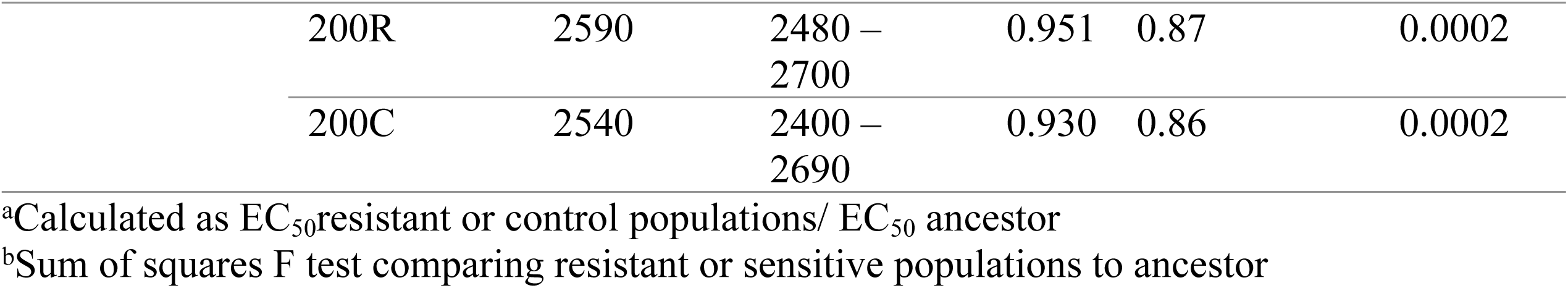
Mean EC_50_ values of evolved ivermectin resistant lines to the cyclooctadepsipeptide emodepside and benzimidazole albendazole as determined by larval development assays.

| **Anthelmintic** | **Population** | **$EC_{50}$**<br>**[nM]** | **95% CI** | **$R^2$** | **Fold**<br>**change<sup>a</sup>** | **P**<br>**value<sup>b</sup>** |
| --- | --- | --- | --- | --- | --- | --- |
| **Emodepside** | Ancestor | 18.27 | 17.50 – 19.07 | 0.927 | 1.00 |  |
|  | 2000R | 91.91 | 88.71 – 95.22 | 0.932 | 5.03 | < 0.0001 |
|  | 2000C | 23.95 | 22.93 – 25.01 | 0.925 | 1.31 | < 0.0001 |
|  | 1000R | 103.4 | 99.04 – 107.9 | 0.912 | 5.65 | < 0.0001 |
|  | 1000C | 16.38 | 15.68 – 17.11 | 0.912 | 0.89 | 0.0006 |
|  | 200R | 97.49 | 94.29 – 100.8 | 0.950 | 5.33 | < 0.0001 |
|  | 200C | 18.68 | 17.95 – 19.44 | 0.922 | 1.02 | 0.4573 |
| **Albendazole** | Ancestor | 2950 | 2790 – 3120 | 0.934 | 1.00 |  |
|  | 2000R | 3190 | 3030 – 3350 | 0.936 | 1.08 | 0.04 |
|  | 2000C | 3410 | 3020 – 3850 | 0.836 | 1.15 | 0.02 |
|  | 1000R | 3300 | 3160 – 3450 | 0.952 | 1.11 | 0.002 |
|  | 1000C | 2730 | 2520 – 2950 | 0.895 | 0.92 | 0.10 |

|  |  |  |  |  |  |
| --- | --- | --- | --- | --- | --- |
| 200R | 2590 | 2480 – 2700 | 0.951 | 0.87 | 0.0002 |
| 200C | 2540 | 2400 – 2690 | 0.930 | 0.86 | 0.0002 |

<sup>a</sup>Calculated as EC<sub>50</sub>resistant or control populations/ EC<sub>50</sub> ancestor
<sup>b</sup>Sum of squares F test comparing resistant or sensitive populations to ancestor

The ancestor and the endpoint of the IVM-treated and control lines were further tested in larval development assays against other classes of anthelmintics: albendazole (benzimidazole), levamisole (imidazothiazole), and monepantel (amino-acetonitrile). For albendazole, all evolved IVM-treated lines and control lines showed similar responses, with mean EC_50_ values between 2.54 µM and 3.41 µM, to the ancestor (2.95 µM) (Fig. 6D, Table 2). There was no consistent pattern for the IVM-treated lines, which showed changes between 0.87 fold (200R) and 1.11 fold (1000R) compared to the ancestor (Table 2). Although most changes in EC_50_ values were significant, there was no pattern, and effect sizes were very small, suggesting that selection for IVM does not lead to systematic changes in susceptibility to albendazole.

Another class of anthelmintics tested was the imidazothiazoles, of which levamisole was used. All IVM-treated and control lines had mean EC_50_ values ranging between 4.58 µM to 10.78 µM (Fig. 6E). The ancestor had an EC_50_ value of 7.51 µM (Table 3). The 2000 and 1000 IVM-treated lines significantly increased their mean EC_50_ value compared to the ancestor, but this increase was only 1.4 fold. The 200 IVM-treated line had a mean EC_50_ value, which decreased compared to the ancestor’s EC_50_ and was not significantly different. All control lines had lower mean EC_50_ values than their corresponding IVM-treated lines. The mean EC_50_ value of these control lines was also significantly different from the ancestor. Overall, while the 2000 and 1000 IVM-treated lines showed a significant difference from the ancestor, the fold change was not high enough to be considered resistant.

**Table 3.** EC_50_ values of evolved ivermectin resistant populations to anthelmintics interacting with nicotinic acetylcholine receptors levamisole and monepantel determined by larval development assays.

| Anthelmintic | Population | EC<sub>50</sub><br>[nM] | 95% CI | R<sup>2</sup> | Fold<br>Change<sup>a</sup> | P<br>value<sup>b</sup> |
| --- | --- | --- | --- | --- | --- | --- |
| Levamisole | Ancestor | 751 | 697 – 809 | 0.906 | 1.00 |  |
|  | 2000R | 1054 | 967 – 1147 | 0.880 | 1.40 | < 0.0001 |
|  | 2000S | 591 | 550 – 634 | 0.925 | 0.78 | < 0.0001 |
|  | 1000R | 1078 | 1012 – 1149 | 0.920 | 1.43 | < 0.0001 |
|  | 1000S | 541 | 506 – 579 | 0.925 | 0.72 | < 0.0001 |
|  | 200R | 695 | 635 – 760 | 0.878 | 0.92 | 0.192 |
|  | 200S | 458 | 437 – 479 | 0.950 | 0.60 | < 0.0001 |
| **Monepantel** | Ancestor | 10.21 | 9.76 – 10.68 | 0.912 | 1.00 |  |
|  | 2000R | 9.64 | 9.16 – 10.15 | 0.894 | 0.94 | 0.987 |
|  | 2000S | 9.83 | 9.14 – 10.57 | 0.861 | 0.96 | 0.366 |
|  | 1000R | 10.76 | 10.34 – 11.20 | 0.924 | 1.05 | 0.084 |
|  | 1000S | 9.98 | 9.50 – 10.49 | 0.895 | 0.97 | 0.506 |
|  | 200R | 8.68 | 8.28 – 9.10 | 0.897 | 0.85 | < 0.0001 |
|  | 200S | 10.89 | 10.42 – 11.38 | 0.901 | 1.06 | 0.045 |

<sup>a</sup>Calculated as EC<sub>50</sub>resistant or control populations/ EC<sub>50</sub> ancestor
<sup>b</sup>Sum of squares F test comparing resistant or sensitive populations to ancestor

The last anthelmintic tested was monepantel from the group of amino-acetonitrile derivatives. The ancestor had an EC_50_ of 10.21 nM (Table 3). The IVM-treated and control lines responded similarly to the ancestor, with the mean EC_50_ values ranging between 8.68 nM and 10.89 nM (Table 3). No drug response pattern was detectable for the lines (Fig. 6F). The lines that showed a significant difference from the ancestor were the 200 IVM-treated line and its corresponding control line. The 200 IVM-treated line had a lower mean EC_50_ than the ancestor, and the 200 control line had a mean EC_50_ value higher than the ancestor. However, as the fold change was around one, no indication of sensitivity or resistance was observed in these populations.

### Lack of a fitness cost for ivermectin selected populations

The evolution of resistance against specific stressors, including drugs, often comes with a significant fitness cost, especially in the absence of the stressor. Therefore, we assessed whether the evolved populations showed indications of a fitness cost using the larval developmental assay. Worms were allowed to develop in V medium for 48-52 hours for the larval development assay (Fig. 7A). Using a non-parametric one-way ANOVA with the Kruskal-Wallis test, we observed no statistical difference between the lines. Therefore, the statistical significance from the one-way ANOVA test might be due to deviations from normality rather than differences between the median developments of the lines. Hence, no consistent pattern was observed for a possible fitness cost of our evolved IVM-treated lines. Although the distribution of developed worms for the ancestor was greater than for the evolved populations, further studies and more sensitive testing are needed to determine if there is indeed no fitness cost associated with IVM resistance.

**Fig. 7.**
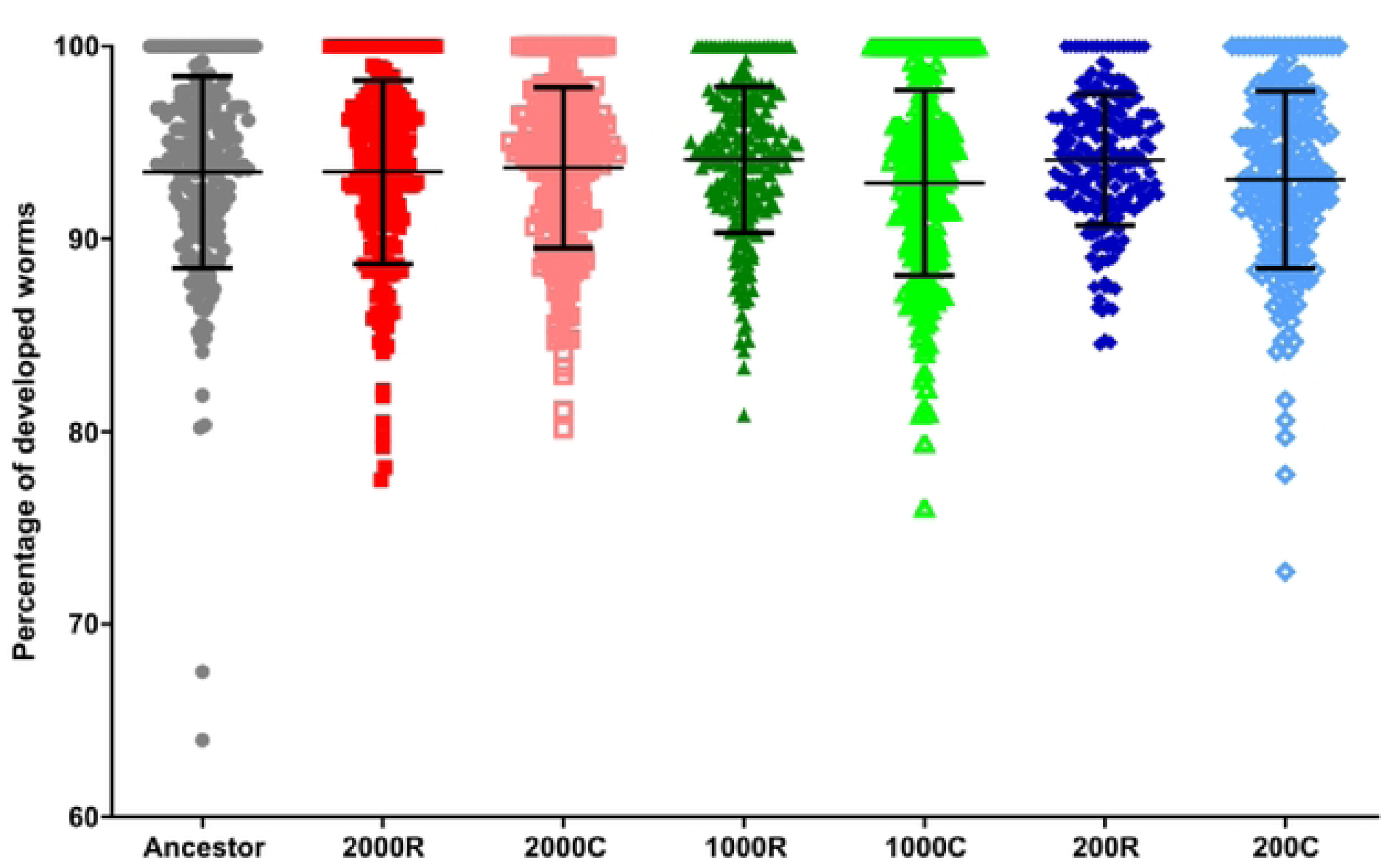
Development of the ancestor and final evolved populations in vehicle controls. Worms were measured for development after a 48 hour incubation at 20 °C in V medium ,vith 0.4% DMSO. Ancestor is shown as black squares, the 2000 IVM-treated line (2000R) as red squares, the 2000 control line (2000C) as salmon squares, the 1000 IVM-treated line (IOOOR) as dark green triangles, the I000 control line (IOOOC) as light green triangles, the 200 IVM-treated line as dark blue diamonds, the 200 control line as light blue diamonds. The percentage of worms that fully developed to the L4/adult stage for individual replicates is graphed as a scatter plot. Median and 95% CI are shown. Kruskal­ Wallis test was performed and no significant differences were found.

## Discussion

The findings presented in this study shed light on the intrinsic factor of population size and its influence on the rate of IVM resistance in an evolution experiment. While various research groups have previously generated IVM resistant worm strains, this study focuses on critical factors that could affect the rate of resistance evolution. Notably, while previous selection experiments typically involved selecting a single line of hermaphrodites, this study examined the effect of population size using twelve biological replicates for each population size. Additionally, an equal number of control lines evolved in the absence of drug selection to monitor the effects of genetic drift and adaptation to the general conditions of the experimental approach. The *in vitro* evolution experiment presented here was further supported by computational modeling, which successfully mirrored the experimental results and provided insights into the potential outcomes of allowing the worms to evolve for an extended period.

We found that population size strongly and significantly affects how quickly a worm population can evolve resistance to IVM. The 2000 IVM-treated line adapted the fastest, reaching a final concentration of 15 nM IVM. In contrast, the 200 IVM-treated line took the longest time to adapt, reaching resistance to only 8 nM IVM (Fig. 2A). This tenfold difference in size resulted in worms being approximately half as adapted. Another factor that could influence the rate of adaptation to IVM among different populations is the genetic variance resulting from male outcrossing. Although we worked with a derivative of the lab wild-type N2 and not a natural isolate, we included males within the population. In an N2 background, males are usually lost within a few generations under standard experimental evolution conditions (45,48–50). Instead, in our evolution experiment, males were consistently maintained at approximately 20% in the presence of IVM and thus at significantly higher levels than under control conditions (Fig. 2B, C). Previous work demonstrated that males are favored in *C. elegans* populations in the presence of environmental stressors (51). For example, males were favored during coevolution of *C. elegans* with bacterial pathogens such as *Serratia marcescens* (46) or *Bacillus thuringiensis* (47), most likely because of a selective advantage of outcrossing under these conditions (47). A similar higher proportion of males was observed in *C. elegans* populations adapting to hyperosmolarity, heat, or chronic irradiation (52–54). We thus conclude that the outcrossing and, as such, the continuous presence of males similarly provided a selective advantage during experimental adaptation to IVM (Fig. 2). Intriguingly, the proportion of males was higher in the more rapidly adapting larger populations, but this could be an artifact due to a stochastic loss of males in the L4/adult stage in the 200 IVM-treated line to count. The higher proportion of males in the larger IVM-treated lines may suggest that the high population size either favors the selection of traits that ensure higher male frequencies or prevents the random loss of males, in both cases promoting the faster adaptation to the environmental stressor.

The evolution of IVM resistance was substantially faster for our mixed-sex population than in previous studies based on populations with only hermaphrodites. In detail, in the study by James and Davey (2009), the hermaphrodite populations required 43 generations to evolve resistance to 6.9 nM IVM and 60 generations to 11.4 nM IVM (26). In comparison, our experiment took 38/39 generations from the introduction of IVM to the final IVM selected populations. In a different study, hermaphrodite populations needed 40 generations to evolve resistance to 5.7 nM MOX (27). These observations further support our conclusion that males and, most likely, their contribution to outcrossing and population genetic variation significantly increase the rate of adaptation to environmental challenges.

Although the evolution experiment yielded similar results to those conducted in the past, the setup was quite different. This was the first IVM evolution experiment that did not utilize nematode growth media (NGM) but instead V medium. As worms exposed to IVM in V medium were previously shown to be more sensitive than worms on NGM (42), it would be interesting to assess the difference between the IVM-treated lines in this study and those previously created to compare their resistance levels. Interestingly, James and Davey also encountered a delay in the rate of resistance evolution to IVM at 6.9 nM, similar to the delay observed in this study for the 2000 and 1000 IVM-treated lines at 6 nM and 5 nM, respectively (26). However, the delay for the 2000 and 1000 IVM-treated lines only lasted for a maximum of 6 generations compared to the 30-generation delay observed previously (26). The reduced delay in adaptation in our study could be due to the presence of males and their effect on outcrossing and genetic variability.

In our study, twelve population replicates within an IVM-treated line were independently subjected to IVM to assess if evolutionary trajectories to resistance vary among individual populations, unlike previous experiments that were based on only a single biological replicate. In our experiment, all 12 population replicates for all three IVM-treated lines were able to adapt to IVM (Fig. 2). Assessment by larval development assays found that all twelve population replicates for the three IVM-treated lines were similar concerning their respective IVM resistance phenotype, and fold changes between the IVM-treated and control lines decreased as the population size decreased (Fig. 6, Table 1). The difference in resistance between different lines was also observed in James and Davey’s study, as their worms resistant to 6.9 nM IVM (IVR6) had an EC_50_ value of 9.0 nM and the EC_50_ value of the 11.4 nM IVM (IVR10) worms was 36.5 nM (26). Despite using different methods for assessing and obtaining IVM-treated lines, the overall trend remains consistent: lines with lower resistance exhibit lower EC_50_ values (26,27). Similar to previous studies, the IVM-treated lines were also resistant to moxidectin. James and Davey reported that IVR6 was less resistant to moxidectin than IVR10 (26), which we also observed when comparing the resistance of the IVM-treated 2000, 1000, and 200 lines.

The IVM-treated lines were also found to be resistant not only to moxidectin but also to emodepside. The observed resistance to moxidectin was expected, as both IVM and MOX affect GluCls, causing hyperpolarization by the influx of chloride ions, leading to paralysis and death (8,15). However, the encountered cross-resistance to emodepside was surprising. Emodepside activates SLO-1 (BK) potassium channels and G-protein-coupled receptors (GPCRs) in nematodes, leading to paralysis and death (55). Emodepside is an anthelmintic active against a broad spectrum of nematodes, including migratory and histotropic stages (56). It has been used successfully to clear infection of multi-resistant (benzimidazoles, levamisole or pyrantel, and macrocyclic lactones) *Haemonchus contortus* in sheep (57) and *Ancylostoma caninum*, canine hookworm, in greyhounds (57,58). Therefore, emodepside exerts its anthelmintic effects on targets that are distinct from the other commonly used anthelmintics. For the human whipworm, *Trichuris trichiura*, emodepside had an 83% cure rate at a 5 mg dose and 100% at 15 mg, compared to 400 mg of albendazole, which had a cure rate of 76% (59). Due to its selective targets and high efficiency, emodepside is currently being considered for treatment as an adulticide against human onchocercosis (including river blindness) and human trichurosis and hookworm disease (55,59). Previously, IVM was the sole drug used to control onchocercosis in Africa since 1995, in the Americas in the 2000s, and in Yemen since 1992 (60). Although the use of IVM to control onchocerciasis led to many areas reporting the elimination of onchocerciasis as a public health problem (61–65), it is only active against the microfilaria and able to sterilize the females for several months. This means that elimination programs require a treatment schedule for decades, with patients being treated once or twice yearly. However, due to the sole use of IVM to treat onchocerciasis, there are growing concerns about resistance and increasing reports of reduced effectivity. Therefore, emodepside, along with other anthelmintics, is being considered as a replacement for or an additional adulticidal treatment arm to IVM in controlling onchocerciasis. However, to the best of our knowledge, our study is the first to report that IVM-resistant *C. elegans* are also cross-resistent to emodepside. This finding raises questions about whether emodepside is the right anthelmintic for the future as a more effective treatment of nematode infections, for cross-resistance may already be present. We suggest that further research should be conducted to assess how widespread similar levels of cross-resistance of ML-resistant parasitic nematodes to emodepside can be found. The findings mentioned above for ML-resistant *H. contortus* and *A. caninum* suggest that the problem might not be relevant immediately in veterinary practice. Still, the results presented here suggest that close monitoring is advised.

Despite the cross-resistance to emodepside, our IVM-resistant populations showed the same trend as observed by James and Davey in relation to the other classes of anthelmintics (26). Our IVM-resistant populations were susceptible to albendazole, similar to their findings. Additionally, there was a small increase in the EC_50_ value for our IVM-resistant populations to levamisole; however, this small change in the EC_50_ value does not indicate the same extent of resistance as seen with moxidectin or emodepside. Furthermore, we also tested monepantel and found that our IVM-resistant populations remained fully susceptible to this anthelmintic.

Our *in vitro* evolution experiment was complemented by polygenic computational modeling using a pharmacodynamic framework. In agreement with experimental observations, the model showed that the largest population had the highest rate of resistance evolution to ivermectin, with remarkable quantitative agreement between the concentrations reached by simulated populations and those reached by the *in vitro* IVM-treated lines. However, the computational and in vitro methods diverged the computational model showed that some populations failed to adapt beyond 2.0 nM of IVM, which was not observed in the *in vitro* experiment. It is assumed that, by chance, the populations in the *in vitro* experiment happened to adapt beyond 2.0 nM. Additionally, if the number of replicates in the *in vitro* experiment matched those in the model, it is likely that some populations would also fail to adapt beyond 2.0 nM.

In the extended model simulating 80 generations, the largest population (2000) remained adapted to a similar concentration of IVM as in the 40 generation model, while the 1000 and 200 population sizes reached higher concentrations of IVM adaptation in the extended model. This suggests that the initial adaptation in the larger population sizes is more likely due to the increased mutation supply. Larger population sizes have been previously linked to faster rates of resistance in evolution experiments and to genetic diversity (66,67). However, as the computational model was set to mimic the *in vitro* experiment and not further investigate the extreme effects of population size. Therefore, the difference in population size might not be significant enough to elicit the same response to IVM resistance over long-term adaptation rates.

The advantage of the computational model is the ability to generate many repeats and to investigate the evolutionary dynamics and genetic composition in full detail. Our analysis revealed that population size significantly influences which resistance mutations become fixed in a population. Smaller populations show higher randomness in gene fixation, while larger populations favor mutations with greater benefits. This suggests that the random occurrence of mutations significantly impacts adaptation trajectories, especially in smaller populations (67–69). In larger populations, subtle differences in the benefits and costs of individual mutations can significantly influence their likelihood of becoming fixed. To thoroughly investigate the fixation of mutations, the computational model was extended from 40 generations to 80. Also, even after 80 generations, none of the populations achieved homozygous mutations on all six loci (Fig. 3 C, D), suggesting that there is additional potential for resistance evolution.

However, the model may differ from the biological reality: While the costs and benefits of individual mutations accumulate, some mutations may epistatically interact in an organism. The model’s assumption of multiplicative fitness costs (see Supplementary material for details) might need to be revisited, especially their relevance under continuous ivermectin exposure. More experimental studies that determine the phenotypic effects of combinations of multiple resistance mutations are needed before the model can be adjusted properly. Previous population genetic modeling studies, for instance, Coffeng et al. (2024) and Schwab et al. (2006), did not assume fitness costs associated with resistance mutations. However, there is insufficient data on the cost of resistance in parasitic worms at the moment. Moreover, how these would combine for mutations at multiple loci is unclear. Obviously, more experimental and modeling research is needed in this area.

## Conclusions

Our study demonstrated that population size and, most likely, the frequency of outcrossing and genetic variation (as approximated by the frequency of males in the present study) impact the rate of IVM resistance evolution. The IVM-treated lines were side-resistant to moxidectin and cross-resistant to emodepside, suggesting that experimental evolution favored broad-range resistance mechanisms.

Further, the computational model, which was set up using key parameters for the *in vitro* evolution experiment, showed very similar results, showing that population size is a crucial parameter. Additional modeling also suggested that the effects of population size might level off to some extent in the long term, which indicates that population size is more important for the speed of resistance evolution than for the outcome after a high enough number of generations. The additional insight provided by the model suggests that while limited mutational supply plays a critical role in smaller populations, in larger populations, clonal interference leads to the preferential fixation of mutations with larger benefits and smaller fitness costs.

## Acknowledgements

We would like to thank the Caenorhabditis Genetics Center for the worms provided which is funded by the NIH Office of Research Infrastructure Programs (P40 OD010440) and to Agnes Piecyk for the initial discovery of Viscous medium as a *C. elegans* medium source.

## Author Contributions

Jacqueline Hellinga completed all of the *C. elegans* experiments and contributed to the analysis and writing of the manuscript.

Barbora Trubenova performed the computational modeling and contributed to the writing of the manuscript.

Jessica Wagner helped with the *in vitro* evolution *C. elegans* experiment.

Roland R. Regoes helped with the computational modeling and the revisions of the manuscript.

Jürgen Krücken helped with the setup of the *in vitro* evolution *C. elegans* experiment, statistical analysis of the data, and revisions of the manuscript.

Hinrich Schulenburg helped with the setup of the *in vitro* evolution *C. elegans* experiment, provided the knowledge about Viscous medium, analysis of the computational modeling and revisions of the manuscript.

Georg von Samson-Himmelstjerna helped with the setup of the *in vitro* evolution *C. elegans* experiment, analysis of the data, and revisions of the manuscript.

## Supporting Information captions

**Fig. S1**. Assessment of progeny production of male induced C. elegans WMB1133 strain population after treatment with ethylmethanesulfonate (EMS). 15 hermaphrodites from untreated and EMS treated (F1 generation) were allowed to lay eggs for 16 hours with 1 worm on each NGM plate. The eggs then were incubated for 96 hours at 16C and the number of alive worms on each plate were counted. The average is graphed and shown is the average with the top and bottom bar indicating the standard deviation.

**Fig. S2**. Parallel exposure of IVM-treated 2000 population from generation 3 to 6 for the working (orange background) and challenger (yellow background) lines. Red X indicates the population did not survive the IVM concentration.

**Fig. S3**. Worm growth during the in vitro evolutionary experiment in comparison to the starting population size. Dashed line indicates the starting population size, the dotted line indicates the population size 10X higher than the starting population size. Pink lines indicate when the population was transferred to a higher concentration of IVM. A) 2000 population, B) 1000 population, C) 200 population.

**Fig. S4.** Fitness of individual strains. A) Pharmacodynamic curves of various sub-populations as a function of genotype and drug concentration. The highest fitness of 1 belongs to the wild type in the absence of drugs, and EC50 of the wild type is set to 1. Different colors represent the number of loci that carry two mutated alleles. Benefit and cost values are shown in Table S2. B-D) The fitness of individual genotypes at concentrations 0, 2 and 10.

**Table S1.** Increase resistance of C. elegans against ivermectin.

**Table S2**. Mutational effects used in the simulations

**Table S3.** Sequence of ivermectin concentrations. Simulation started with no ivermectin. If the population of worms was deemed sufficiently adapted, concentration was increased. S2.2

**Fig. S5.** Distribution of the final concentration reached after 40 generations. Horizontal lines represent medians. The final genetic compositions of the populations at generation 40 differed profoundly among different simulation trials. While the average number of homozygous mutants increased over time in all three sample sizes (Figure 4 B), in none of the simulations were full mutants observed, suggesting that the populations could evolve further.

**Figure S6.** Genetic composition of 15 randomly selected final populations of 200 worms after 40 generations of evolution. On the X axis, different loci are portrayed, and on the Y axis, the genotypes that are possible for each locus. The rectangle’s color corresponds to the fraction of individuals having a particular genotype on that particular locus. Values in each column add to 1.

**Figure S7.** Genetic composition of 15 randomly selected final populations of 1000 worms after 40 generations of evolution. On the X axis, different loci are portrayed, and on the Y axis, the genotypes that are possible for each locus. The rectangle’s color corresponds to the fraction of individuals having a particular genotype on that particular locus. Values in each column add to 1.

**Figure S8.** Genetic composition of 15 randomly selected final populations of 2000 worms after 40 generations of evolution. On the X axis, different loci are portrayed, and on the Y axis, the genotypes that are possible for each locus. The rectangle’s color corresponds to the fraction of individuals having a particular genotype on that particular locus. Values in each column add to 1.

**Figure S9.** Concentration at which the worm population grew as a function of time. Blue solid lines correspond to individual simulations; the red dashed line represents the mean. Population size (A) 200, B) 1000, C) 2000. **Figure S10.** Distribution of the final concentration reached after 40 generations. Horizontal lines represent medians.

**Figure S11.** Simulations of the neutral evolution. The average number of homozygous mutations per worm present within the population at a certain time point: 200 (blue), 1000 (green) and red (2000).

**Figure S12**. The average genetic composition of 100 final populations: (A) 200 (B) 1000 (C) 2000. On the X axis, different loci are portrayed, and on the Y axis, the genotypes that are possible for each locus. The rectangle’s color corresponds to the fraction of individuals having a particular genotype on that particular locus. Values in each column add to1.

